# Hormonal stimulation induces broader decidualization responses than cAMP alone in 3D human endometrial organoids

**DOI:** 10.64898/2026.03.25.714293

**Authors:** Shichao Liu, Jiyang Zhang, Tingjie Zhan, Qiang Zhang, Nataki C. Douglas, Xiaoqin Ye, Shuo Xiao

## Abstract

The human endometrium undergoes cyclic, hormone-driven remodeling that establishes a transient window of receptivity required for embryo implantation, placentation, and maintenance of pregnancy. Decidualization of endometrial stromal cells is a central component of this process and can be induced *in vitro* using cAMP alone or in combination with ovarian steroid hormones (EPC: estradiol, progesterone, and cAMP). Although cAMP activates the core decidual transcriptional program, whether hormone supplementation induces a more physiologically relevant response remains unclear, particularly in 3D endometrial organoid (Endo-organoid) models which have emerged as a new alternative methodology (NAM). Here, we compared morphological and transcriptomic responses of human endometrial stromal cell-derived Endo-organoids undergoing decidualization induced by cAMP or EPC stimulation. EPC-treated Endo-organoids exhibited enhanced structural remodeling and more advanced morphological transformation compared with cAMP-treated organoids. RNA-seq analysis revealed substantial overlap in canonical decidual gene expression between the two conditions, but EPC induced broader transcriptional and pathway-level changes, including enrichment of metabolic, stress-response, and differentiation-related processes. Together, these findings demonstrate that while cAMP activates the core decidual program, EPC elicits a broader and more physiologically relevant decidualization response in 3D human Endo-organoids, providing guidance for optimizing Endo-organoids to study endometrial receptivity, implantation, and early pregnancy success.

---

The endometrium undergoes cyclic, hormone-driven remodeling that establishes a transient window of receptivity necessary for successful embryo implantation, placentation, and maintenance of pregnancy. During this process, endometrial stromal cells undergo decidualization, in which progesterone secreted by the ovarian corpus luteum differentiates them from fibroblast-like cells into rounded decidual cells, accompanied by profound functional transformation, creating an immunologically tolerant uterine environment to support embryo implantation and subsequent embryonic and fetal development. Due to ethical concerns of including human subjects, there are few rigorous, scalable approaches for studying endometrial receptivity. *In vitro* culture of three-dimensional (3D) endometrial organoids (Endo-organoids) has recently emerged as a powerful platform for modeling endometrial transformation [1-5]. Endo-organoids preserve essential characteristics of the endometrial microenvironment, including 3D spatial architecture, cell-cell communication, and extracellular matrix (ECM), which cannot be recapitulated in conventional two-dimensional (2D) cultures.

Several protocols have been employed to induce *in vitro* decidualization, including stimulation with cyclic AMP (cAMP) alone and the EPC protocol consisting of estradiol, medroxyprogesterone acetate (MPA, a synthetic progestin), and cAMP [6]. Accumulating evidence indicates that the elevation of intracellular cAMP in endometrial stromal cells represents a key molecular initiating event in decidualization and functions as a central mediator enabling progesterone signaling to activate decidua-specific transcriptional networks. The rise of intracellular cAMP levels in stromal cells might be controlled by autocrine, paracrine, or endocrine factors, including relaxin (RLN), corticotropin-releasing hormone (CRH), and/or prostaglandin E2 (PGE2) [6, 7]. Although cAMP efficiently induces the core decidual transcriptional program, it remains unclear whether and how the supplementation with ovarian steroid hormones produces a more physiologically relevant decidual response. Moreover, how 3D Endo-organoids respond to different decidualization stimuli remains unknown, which limits our ability to optimize Endo-organoid models and to interpret their functional relevance for studying endometrial receptivity. To address these questions, we compared the transcriptomic responses of human endometrial stromal cell-derived Endo-organoid undergoing *in vitro* decidualization induced by cAMP or EPC stimulation.

Human endometrial stromal cells were seeded into agarose-based 3D Petri Dish molds to generate Endo-organoids, followed by six days of decidualization induction with 0.5 mM cAMP alone or EPC (10 nM E2, 1 μM MPA, and 0.5 mM cAMP). Bright-field imaging revealed that vehicle-treated Endo-organoids remained compact and slightly decreased in size during culture (0.64 ± 0.13, n = 28); endo-organoids treated with cAMP displayed modest structural changes with a size similar to day 0 (1.04 ± 0.14, n = 21); in contrast, EPC-treated Endo-organoids significantly expanded (1.25 ± 0.38, n = 24) compared with vehicle- and cAMP-treated organoids (Figure 1A-1B). Cytoskeletal organization was examined by F-actin immunofluorescent staining (Figure 1C). Control Endo-organoids displayed uniformly spindle-shaped stromal cells, indicative of a pre-decidualization phenotype. cAMP induced modest structural change, with most stromal cells in the organoid core retaining spindle morphology and only a small fraction of peripheral cells exhibiting partial rounding. In contrast, EPC-treated organoids showed more extensive decidual transformation, with interior cells adopting a rounded morphology and surface cells becoming nearly completely rounded.

**Figure 1.**
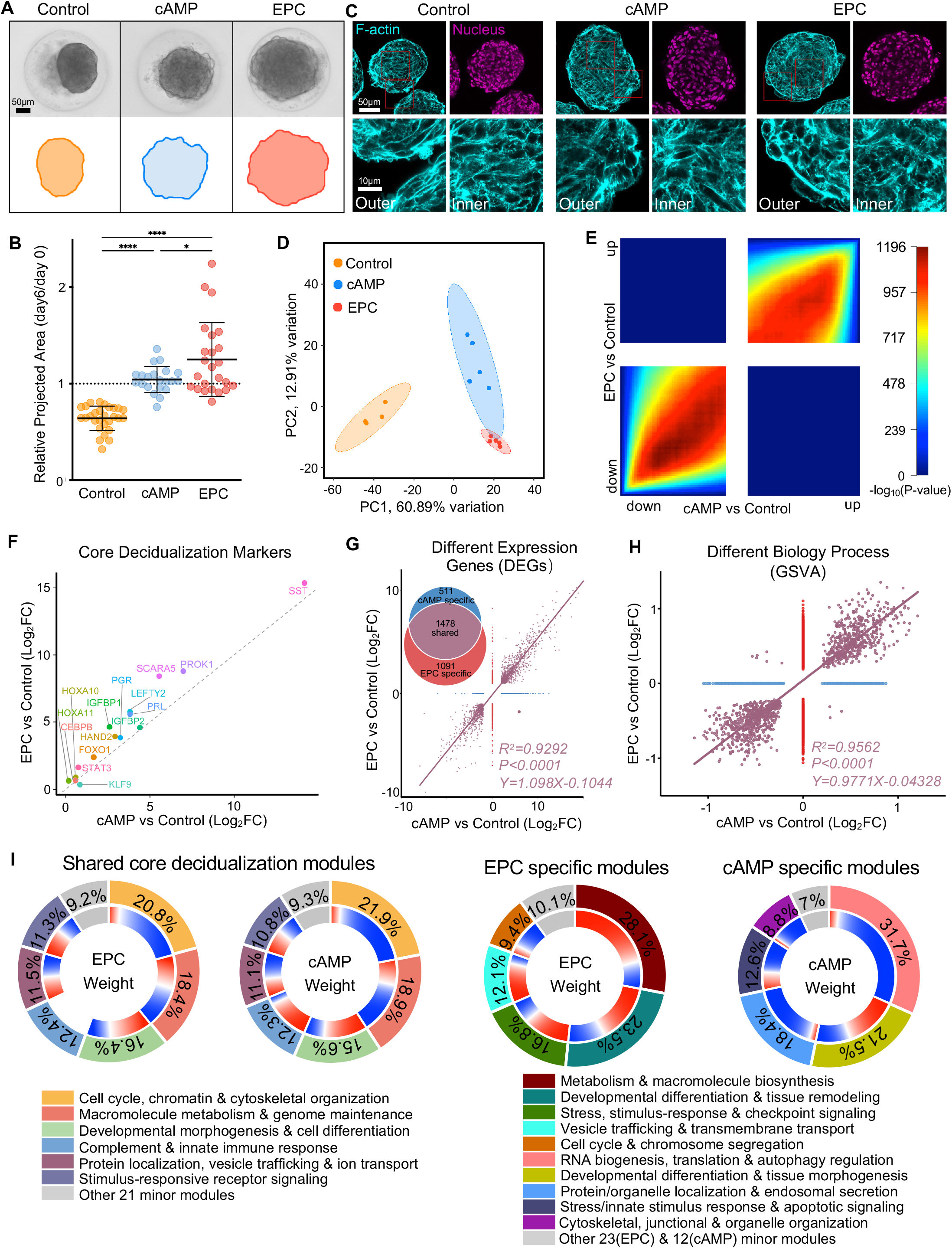
Morphological and transcriptomic characterization of decidualization responses in 3D human endometrial organoids (Endo-organoids) induced by cAMP or EPC. **(A)** Bright-field images of Endo-organoids after 6 days of culture and treatment with vehicle (Control), cAMP, or EPC, together with corresponding projected areas. Scale bars are shown as indicated. **(B)** Quantification of Endo-organoid growth represented by the ratio of projected area (day 6 / day 0). One-way ANOVA was performed (\*\*\*\**P < 0*.*0001*, \**P < 0*.*05*). n = 28, 21, and 24 organoids in Control), cAMP), and EPC groups, respectively. **(C)** F-actin immunofluorescence staining of Endo-organoids at day 6. Cyan indicates F-actin (phalloidin), and magenta indicates nuclei (DAPI). The “Inner” and “Outer” panels correspond to the regions highlighted in red boxes in the upper images. Scale bars are shown as indicated. **(D)** Principal component analysis (PCA) of RNA-seq data showing separation among Control, cAMP, and EPC groups. **(E)** Genome-wide concordance analysis using rank–rank hypergeometric overlap (RRHO), demonstrating strong overlap in both upregulated and downregulated gene sets between cAMP and EPC conditions. **(F)** Scatter plot of canonical decidual marker genes showing log_2_ fold changes (EPC vs Control, y-axis; cAMP vs Control, x-axis). The dashed line indicates x = y. All markers were significantly upregulated with adjusted *P < 0*.*05*, and 14 of 15 genes (except KLF9) lie above the diagonal, indicating stronger induction by EPC. **(G)** Classification of differentially expressed genes (DEGs) into shared (n = 1,478), EPC-specific (n = 1,091), and cAMP-specific (n = 511) groups. Linear regression of shared DEGs shows strong concordance between conditions. **(H)** GSVA-based pathway analysis using Gene Ontology (GO) biological process gene sets. Pathways were categorized into shared, EPC-specific, and cAMP-specific groups, and shared pathway enrichment scores were highly correlated between conditions. **(I)** Functional module organization of pathways in shared, EPC-specific, and cAMP-specific groups based on GO semantic similarity. The outer ring represents the weighted contribution of each module, while the inner ring indicates the proportion of activated (red) and suppressed (blue) pathways. A higher red proportion indicates predominant activation, whereas a higher blue proportion indicates predominant suppression within each module.

To characterize molecular responses to decidualization induction at the transcriptomic level, RNA sequencing (RNA-seq) was performed on Endo-organoids collected after 6 days of stimulation. Principal component analysis (PCA) revealed clear separation between control and stimulated groups, indicating substantial transcriptional remodeling induced by cAMP or EPC (Figure 1D). Although cAMP- and EPC-treated organoids clustered closely, they remained distinguishable, suggesting largely shared but divergent transcriptional responses.

Global transcriptomic concordance between the two decidualization stimulations was further examined using rank-rank hypergeometric overlap (RRHO) analysis based on genome-wide differential expression rankings [8]. The RRHO heatmap revealed strong enrichment signals in both the double-upregulated and double-downregulated quadrants, whereas minimal signals were observed in the up-down and down-up regions (Figure 1 E, Supplemental Table S1). These results indicate highly consistent transcriptional responses between EPC and cAMP stimulation. Consistently, canonical decidualization markers, such as IGFBP1, PRL, and FOXO1, were markedly upregulated under both conditions (Figure 1F). The log_2_ fold changes ranged from ∼0.34 to 15.34 under EPC stimulation and from 0.58 to 14.18 under cAMP stimulation, with adjusted *P* values from ∼10^-^□ to <10^−3^□. Notably, most decidualization markers exhibited slightly higher induction under EPC stimulation.

We identified 1,478 shared differentially expressed genes (DEGs) together with 1,091 EPC-specific and 511 cAMP-specific DEGs (Supplemental Figure S1 and Supplemental Table S2). Linear regression analysis of shared DEGs demonstrated strong concordance between EPC and cAMP responses (R^2^= 0.929, P < 0.0001), while the substantial numbers of stimulus-specific genes suggested additional transcriptional programs activated by each decidualization stimulation condition (Figure 1G).

To investigate overlapping and unique DEGs at the pathway level without introducing bias from preselected gene subsets, we applied gene set variation analysis (GSVA) using Gene Ontology (GO) Biological Process gene sets [9]. There were 894 shared pathways together with 1124 EPC-specific and 586 cAMP-specific pathways. Enrichment scores of shared pathways also showed strong concordance between EPC and cAMP stimulation (R^2^=0.9562, *P<0*.*0001*, Figure 1H, Supplemental Figure S2, and Supplemental Table S3).

To summarize the functional landscape, pathways were grouped into modules based on GO semantic similarity. Across the three groups, thousands of pathways converged into a limited number of modules, indicating strong functional coordination: 6/27 modules accounted for 90.8% and 90.7% of total weighted pathway activity in EPC and cAMP within the shared pathway set; 5/28 modules accounted for 89.9% of EPC-specific pathways; and 5/17 modules accounted for 93.0% of cAMP-specific pathways (Figure 1I and Supplemental Table S4). Within the shared pathway set, EPC and cAMP displayed nearly identical module distributions and activation patterns. The dominant modules corresponded to cell cycle and cytoskeletal organization (EPC 20.8% vs cAMP 21.9%), macromolecule metabolism (18.4% vs 18.9%), developmental morphogenesis (16.4% vs 15.6%), complement/immune signaling (12.4% vs 12.3%), vesicle trafficking (11.5% vs 11.1%), and stimulus-responsive signaling (11.3 vs 11.8%), which are consistent with key biological processes underlying stromal decidualization *in vivo* [6, 10]. EPC-specific pathways were primarily concentrated in modules related to metabolic adaptation, stress-responsive signaling, developmental differentiation, vesicle trafficking, and cell cycle regulation, indicating broader physiological remodeling with the addition of estradiol and progesterone. In contrast, cAMP-specific pathways were mainly enriched in intracellular process modules associated with RNA biogenesis, transcriptional regulation, intracellular trafficking, and stress-response signaling, many showing predominantly suppressed activity.

Together, our study demonstrates that while cAMP efficiently activates the core decidual transcriptional program, EPC stimulation elicits broader and more physiologically relevant decidual responses in 3D human Endo-organoids. These results provide mechanistic insights into how cAMP and ovarian-derived steroid hormones coordinate endometrial decidualization and provide evidence for optimizing *in vitro* 3D Endo-organoid culture and decidualization induction for studying endometrial receptivity, embryo implantation, and placentation.

## Supporting information

Supplemental Table S1. RRHO Analysis

Supplemental Table S2. DEGs Analysis

Supplemental Table S3. GSVA Analysis

Supplemental Table S4. Modules Clustering Analysis

## Data availability

All associated RNA-seq data, including fastq files and processed raw counts, are available at the Gene Expression Omnibus (GSE325553).

## Author Contributions

S. Liu, T. Zhan, and J. Zhang contributed to experimental design, data collection and analysis, and manuscript writing; Q. Zhang, NC. Douglas, and X. Ye contributed to data interpretation and manuscript editing; S. Xiao conceived the project, contributed to experimental design, manuscript writing and editing, data analysis and interpretation, and provided final approval of the manuscript.

