## Supplemental information for "Hormonal stimulation induces broader decidualization responses than cAMP alone in 3D human endometrial organoids"

[1]Department of Pharmacology and Toxicology, Ernest Mario School of Pharmacy, Rutgers University, Piscataway, NJ 08854, USA; [2]Environmental and Occupational Health Sciences Institute (EOHSI), Rutgers University, Piscataway, NJ 08854, USA; [3]Gangarosa Department of Environmental Health, Rollins School of Public Health, Emory University, Atlanta, GA 30322, USA; [4]Department of Obstetrics, Gynecology and Reproductive Health, New Jersey Medical School (NJMS), Rutgers University, Newark, NJ 07103, USA; [5]Department of Physiology and Pharmacology, College of Veterinary Medicine, University of Georgia, Athens, GA 30602, USA

### Supplementary Methods

#### Primary human endometrial stromal cells

Primary human endometrial stromal cells were generously provided by the laboratory of Dr. Nataki C. Douglas (Center for Immunity & Inflammation; Department of Obstetrics, Gynecology and Women's Health, Rutgers University). Cells were previously isolated and characterized from human endometrial tissues under institutional review board (IRB)-approved protocols at the originating institution, with informed consent obtained from all donors. Cells were transferred in de-identified form.

#### Cell culture and 3D endometrial organoid generation

Primary human endometrial stromal cells were cultured in DMEM/F12 medium (phenol red-free; Gibco, 11039-021) supplemented with 10% charcoal-stripped fetal bovine serum (Thermo Fisher Scientific, A3382101), 10 ng/mL basic fibroblast growth factor (bFGF; Sigma, GF003), and 1% penicillin/streptomycin under standard culture conditions (37°C, 5% $CO_2$).

For organoid formation, $2 \times 10^5$ cells were resuspended in 50 µL MammoCult™ medium (STEMCELL Technologies) and seeded into agarose-based 3D Petri Dish® micromold inserts (MicroTissues Inc.) placed in 24-well plates. An additional 250 µL MammoCult™ medium was added per well to maintain hydration. Cells were allowed to self-aggregate for 24 h to form uniform endometrial organoids.

Following organoid formation, medium was replaced with culture medium. Decidualization was induced under the following conditions: Control: vehicle; cAMP: 0.5 mM cAMP; EPC: 10 nM estradiol (E2), 1 µM medroxyprogesterone acetate (MPA), and 0.5 mM cAMP. Medium was refreshed every 2 days, and organoids were cultured for 6 days.

#### Relative Projected Area Analysis

Bright-field imaging of Endo-organoids was performed at the start of decidualization induction (Day 0) and after 6 days of treatment (Day 6) using an EVOS™ digital inverted microscope (Thermo Fisher Scientific). For each condition, the entire plate was imaged and individual wells were recorded and indexed to ensure that the same organoids were tracked and analyzed between Day 0 and Day 6.

To quantify morphological changes, the projected area of individual organoids was measured using ImageJ software (NIH). For each organoid, boundaries were manually outlined in bright-field images, and the enclosed area was recorded. Relative size change was calculated as the ratio of projected area at Day 6 to that at Day 0 (Day6/Day0).

#### Immunofluorescence staining

Organoids were fixed in 4% paraformaldehyde at room temperature for 1 h and permeabilized in 1% Triton X-100 in PBS at 4°C overnight. Samples were blocked in 1% BSA in PBS for 2 h at room temperature and incubated with rhodamine-conjugated phalloidin (Invitrogen, Cat# R415) for F-actin staining. Nuclei were counterstained using ProLong™ Gold Antifade Mountant with DAPI (Invitrogen, Cat# P36931). Images were acquired using a Leica SP8 confocal microscope.

#### RNA sequencing (RNA-seq) analysis

Total RNA was extracted from organoids after 6 days of treatment using the PicoPure RNA Isolation Kit (Thermo Fisher Scientific) according to the manufacturer's instructions. RNA integrity and concentration were evaluated prior to library construction. Bulk RNA sequencing libraries were prepared and sequenced on an Illumina platform by Novogene Co., Ltd. (Sacramento, CA). Clean reads were aligned to the human reference genome, and gene expression levels were quantified as transcripts per million (TPM) for downstream analyses.

#### Rank–rank hypergeometric overlap (RRHO)

Rank–rank hypergeometric overlap (RRHO) analysis was performed using the RRHO2 R package. Genes from EPC vs control and cAMP vs control comparisons were ranked based on a signed significance score calculated as *−log10(adjusted P value) × sign(log2 fold change)* (supplemental Table 1). Adjusted P values equal to zero were set to 1e−300 prior to transformation. Only genes present in both datasets were retained for analysis. The RRHO matrix was computed to evaluate gene overlap across all rank thresholds using hypergeometric testing (Supplemental Table 1). Heatmaps display −log10 adjusted P values, where higher values indicate stronger enrichment of overlapping gene sets.

#### Differential gene expression analysis

Differential expression analysis between EPC vs control and cAMP vs control was performed using the DESeq2 R package. Genes with log2FoldChange>1 or <-1 and adjusted P value < 0.05 were considered differentially expressed (Supplemental Table 2). Shared and stimulus-specific DEGs were identified by intersection and set difference between the two comparisons.

#### Gene set variation analysis (GSVA)

GSVA was performed using the GSVA R package to estimate pathway enrichment scores for Gene Ontology (GO) Biological Process gene sets. GSVA transforms gene expression data into sample-wise pathway enrichment scores in a non-parametric manner based on the relative expression ranks of genes within each sample. Differential pathway activity between conditions was assessed using limma, and log2 fold changes (logFC) represent differences in GSVA enrichment scores between groups (Supplemental Table 3).

#### Module clustering based on GO semantic similarity

GO pathways were grouped into functional modules using semantic similarity-based clustering implemented in the simplifyEnrichment package. Pairwise GO term similarity was calculated using the Wang method, and clustering was performed using the binary cut algorithm (Supplemental Table 4).

**Module weight and activity quantification**

For each module, pathway weights were calculated as the sum of absolute enrichment scores (|log2FC|) across all pathways within the module (Supplemental Table 4). The relative module weight contribution was defined as:

Module weight (%) = $\Sigma|NES|$ (module) / $\Sigma|NES|$ (all modules)

To evaluate regulatory direction within each module, activation and suppression proportions were calculated as:

Activation proportion = $\Sigma(NES > 0) / \Sigma|NES|$

Suppression proportion = $\Sigma(NES < 0) / \Sigma|NES|$

All statistical analyses were performed in R.

### Supplementary Figures

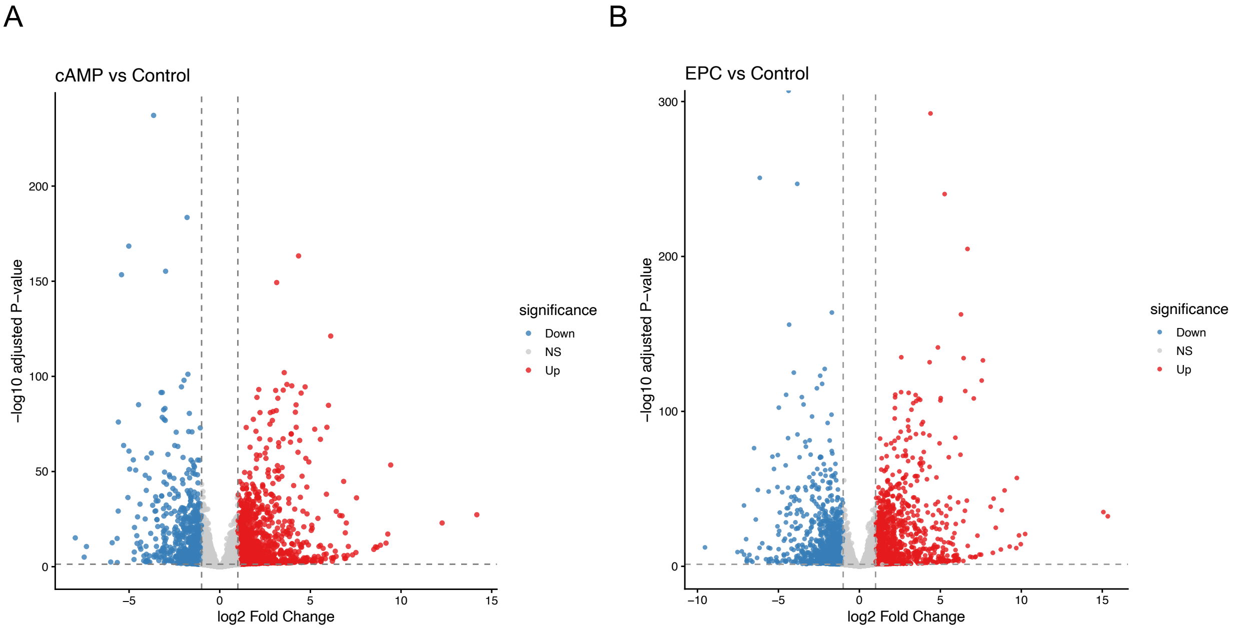


**Supplemental Figure S1. Differential gene expression induced by cAMP and EPC stimulation in 3D Endo-organoids (A–B)** Volcano plots showing differential gene expression in Endo-organoids after 6 days of stimulation. **(A) cAMP vs control; (B) EPC vs control.** Genes were colored based on significance and direction of change: significantly upregulated genes (adjusted P < 0.05 and $\log_2$ fold change > 1) are shown in red, significantly downregulated genes (adjusted P < 0.05 and $\log_2$ fold change < -1) in blue, and non-significant genes in gray. A total of 1,245 upregulated and 744 downregulated genes were identified in the cAMP vs control comparison, whereas 1,166 upregulated and 1,403 downregulated genes were identified in the EPC vs control comparison.

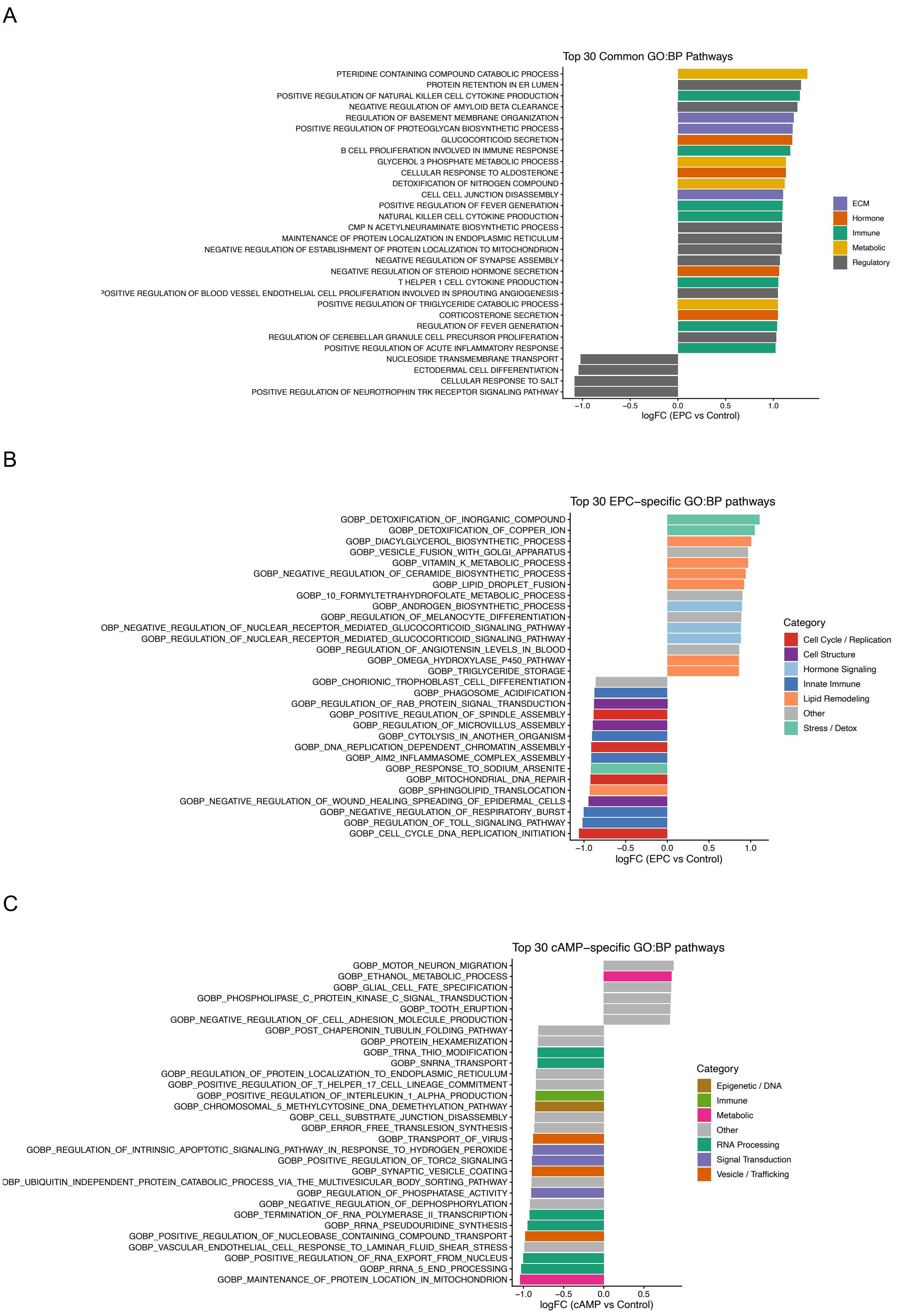
A
Top 30 Common GO:BP Pathways
PTERIDINE CONTAINING COMPOUND CATABOLIC PROCESS
PROTEIN RETENTION IN ER LUMEN
POSITIVE REGULATION OF NATURAL KILLER CELL CYTOKINE PRODUCTION
NEGATIVE REGULATION OF AMYLOID BETA CLEARANCE
REGULATION OF BASEMENT MEMBRANE ORGANIZATION
POSITIVE REGULATION OF PROTEOGLYCAN BIOSYNTHETIC PROCESS
GLUCOCORTICOID SECRETION
B CELL PROLIFERATION INVOLVED IN IMMUNE RESPONSE
GLYCEROL 3 PHOSPHATE METABOLIC PROCESS
CELLULAR RESPONSE TO ALDOSTERONE
DETOXIFICATION OF NITROGEN COMPOUND
CELL CELL JUNCTION DISASSEMBLY
POSITIVE REGULATION OF FEVER GENERATION
NATURAL KILLER CELL CYTOKINE PRODUCTION
CMP N ACETYLNEURAMINATE BIOSYNTHETIC PROCESS
MAINTENANCE OF PROTEIN LOCALIZATION IN ENDOPLASMIC RETICULUM
NEGATIVE REGULATION OF ESTABLISHMENT OF PROTEIN LOCALIZATION TO MITOCHONDRION
NEGATIVE REGULATION OF SYNAPSE ASSEMBLY
NEGATIVE REGULATION OF STEROID HORMONE SECRETION
T HELPER 1 CELL CYTOKINE PRODUCTION
OSITIVE REGULATION OF BLOOD VESSEL ENDOTHELIAL CELL PROLIFERATION INVOLVED IN SPROUTING ANGIOGENESIS
POSITIVE REGULATION OF TRIGLYCERIDE CATABOLIC PROCESS
CORTICOSTERONE SECRETION
REGULATION OF FEVER GENERATION
REGULATION OF CEREBELLAR GRANULE CELL PRECURSOR PROLIFERATION
POSITIVE REGULATION OF ACUTE INFLAMMATORY RESPONSE
NUCLEOSIDE TRANSMEMBRANE TRANSPORT
ECTODERMAL CELL DIFFERENTIATION
CELLULAR RESPONSE TO SALT
POSITIVE REGULATION OF NEUROTROPHIN TRK RECEPTOR SIGNALING PATHWAY
−1.0 −0.5 0.0 0.5 1.0
logFC (EPC vs Control)
ECM
Hormone
Immune
Metabolic
Regulatory
B
Top 30 EPC−specific GO:BP pathways
GOBP_DETOXIFICATION_OF_INORGANIC_COMPOUND
GOBP_DETOXIFICATION_OF_COPPER_ION
GOBP_DIACYLGLYCEROL_BIOSYNTHETIC_PROCESS
GOBP_VESICLE_FUSION_WITH_GOLGI_APPARATUS
GOBP_VITAMIN_K_METABOLIC_PROCESS
GOBP_NEGATIVE_REGULATION_OF_CERAMIDE_BIOSYNTHETIC_PROCESS
GOBP_LIPID_DROPLET_FUSION
GOBP_10_FORMYLTETRAHYDROFOLATE_METABOLIC_PROCESS
GOBP_ANDROGEN_BIOSYNTHETIC_PROCESS
GOBP_REGULATION_OF_MELANOCYTE_DIFFERENTIATION
OBP_NEGATIVE_REGULATION_OF_NUCLEAR_RECEPTOR_MEDIATED_GLUCOCORTICOID_SIGNALING_PATHWAY
GOBP_REGULATION_OF_NUCLEAR_RECEPTOR_MEDIATED_GLUCOCORTICOID_SIGNALING_PATHWAY
GOBP_REGULATION_OF_ANGIOTENSIN_LEVELS_IN_BLOOD
GOBP_OMEGA_HYDROXYLASE_P450_PATHWAY
GOBP_TRIGLYCERIDE_STORAGE
GOBP_CHORIONIC_TROPHOBLAST_CELL_DIFFERENTIATION
GOBP_PHAGOSOME_ACIDIFICATION
GOBP_REGULATION_OF_RAB_PROTEIN_SIGNAL_TRANSDUCTION
GOBP_POSITIVE_REGULATION_OF_SPINDLE_ASSEMBLY
GOBP_REGULATION_OF_MICROVILLUS_ASSEMBLY
GOBP_CYTOLYSIS_IN_ANOTHER_ORGANISM
GOBP_DNA_REPLICATION_DEPENDENT_CHROMATIN_ASSEMBLY
GOBP_AIM2_INFLAMMASOME_COMPLEX_ASSEMBLY
GOBP_RESPONSE_TO_SODIUM_ARSENITE
GOBP_MITOCHONDRIAL_DNA_REPAIR
GOBP_SPHINGOLIPID_TRANSLOCATION
GOBP_NEGATIVE_REGULATION_OF_WOUND_HEALING_SPREADING_OF_EPIDERMAL_CELLS
GOBP_NEGATIVE_REGULATION_OF_RESPIRATORY_BURST
GOBP_REGULATION_OF_TOLL_SIGNALING_PATHWAY
GOBP_CELL_CYCLE_DNA_REPLICATION_INITIATION
−1.0 −0.5 0.0 0.5 1.0
logFC (EPC vs Control)
Category
Cell Cycle / Replication
Cell Structure
Hormone Signaling
Innate Immune
Lipid Remodeling
Other
Stress / Detox
C
Top 30 cAMP−specific GO:BP pathways
GOBP_MOTOR_NEURON_MIGRATION
GOBP_ETHANOL_METABOLIC_PROCESS
GOBP_GLIAL_CELL_FATE_SPECIFICATION
GOBP_PHOSPHOLIPASE_C_PROTEIN_KINASE_C_SIGNAL_TRANSDUCTION
GOBP_TOOTH_ERUPTION
GOBP_NEGATIVE_REGULATION_OF_CELL_ADHESION_MOLECULE_PRODUCTION
GOBP_POST_CHAPERONIN_TUBULIN_FOLDING_PATHWAY
GOBP_PROTEIN_HEXAMERIZATION
GOBP_TRNA_THIO_MODIFICATION
GOBP_SNRNA_TRANSPORT
GOBP_REGULATION_OF_PROTEIN_LOCALIZATION_TO_ENDOPLASMIC_RETICULUM
GOBP_POSITIVE_REGULATION_OF_T_HELPER_17_CELL_LINEAGE_COMMITMENT
GOBP_POSITIVE_REGULATION_OF_INTERLEUKIN_1_ALPHA_PRODUCTION
GOBP_CHROMOSOMAL_5_METHYLCYTOSINE_DNA_DEMETHYLATION_PATHWAY
GOBP_CELL_SUBSTRATE_JUNCTION_DISASSEMBLY
GOBP_ERROR_FREE_TRANSLESION_SYNTHESIS
GOBP_TRANSPORT_OF_VIRUS
GOBP_REGULATION_OF_INTRINSIC_APOPTOTIC_SIGNALING_PATHWAY_IN_RESPONSE_TO_HYDROGEN_PEROXIDE
GOBP_POSITIVE_REGULATION_OF_TORC2_SIGNALING
GOBP_SYNAPTIC_VESICLE_COATING
OBP_UBIQUITIN_INDEPENDENT_PROTEIN_CATABOLIC_PROCESS_VIA_THE_MULTIVESICULAR_BODY_SORTING_PATHWAY
GOBP_REGULATION_OF_PHOSPHATASE_ACTIVITY
GOBP_NEGATIVE_REGULATION_OF_DEPHOSPHORYLATION
GOBP_TERMINATION_OF_RNA_POLYMERASE_II_TRANSCRIPTION
GOBP_RRNA_PSEUDOURIDINE_SYNTHESIS
GOBP_POSITIVE_REGULATION_OF_NUCLEOBASE_CONTAINING_COMPOUND_TRANSPORT
GOBP_VASCULAR_ENDOTHELIAL_CELL_RESPONSE_TO_LAMINAR_FLUID_SHEAR_STRESS
GOBP_POSITIVE_REGULATION_OF_RNA_EXPORT_FROM_NUCLEUS
GOBP_RRNA_5_END_PROCESSING
GOBP_MAINTENANCE_OF_PROTEIN_LOCATION_IN_MITOCHONDRION
−1.0 −0.5 0.0 0.5
logFC (cAMP vs Control)
Category
Epigenetic / DNA
Immune
Metabolic
Other
RNA Processing
Signal Transduction
Vesicle / Trafficking

**Supplemental Figure S2. Functional grouping of top 30 enriched pathways across shared and stimulus-specific responses (A–C)** Top 30 Gene Ontology Biological Process (GO-BP) pathways identified by gene set variation analysis (GSVA) in **(A) shared, (B) EPC-specific, and (C) cAMP-specific** pathway sets. Pathways were ranked based on GSVA enrichment scores and grouped into functional categories according to biological similarity. Each pathway is displayed with its corresponding GSVA enrichment score and statistical significance. Shared pathways primarily represent core decidualization-associated processes. EPC-specific top 30 pathways exhibited greater functional convergence, with only 5/30 unassigned pathways and 15/30 displaying positive GSVA activity. In contrast, cAMP-specific top 30 pathways were largely associated with intracellular regulatory processes, with 14/30 pathways not assigned to major functional modules and 24/30 showing negative GSVA activity.
